# Invasive mosquito species Aedes aegypti and Aedes albopictus are competent vectors for Barmah Forest Virus

**DOI:** 10.64898/2026.08.31.748198

**Authors:** Anna Heitmann, Patrick Höller, Unchana Lange, Amrei Mack, Norbert Becker, Jonas Schmidt-Chanasit, Renke Lühken, Stephanie Jansen

**Affiliations:** Bernhard Nocht Institute for Tropical Medicine, Hamburg, Germany; Institute for Dipterology (IfD), Speyer, Germany; Center for Organismal Studies (COS), University of Heidelberg, Heidelberg, Germany; Faculty of Mathematics, Informatics and Natural Sciences, Universität Hamburg, Hamburg, Germany

**Keywords:** Barmah Forest Virus, *Aedes albopictus*, *Aedes aegypti*, vector competence

## Abstract

Barmah Forest Virus (BFV), an arthropod-born virus transmitted by mosquitoes, is of significant public health concern in Australia and regions in the Pacific. Recent climate change and globalization raise the potential for BFV to extend its geographic distribution. Despite the rising importance of BFV, its vector dynamics remain poorly understood, particularly concerning the vector competence of different mosquito species. This study aims to investigate the vector competence of BFV across various mosquito species, beyond those endemic to the Australasian region, especially focusing on global relevant vector species. No transmission was observed for *Culex quinquefasciatus* and *Cx. torrentium* as well as *Anopheles stephensi*. In contrast, both investigated *Aedes* species, *Ae. aegypti* as well as *Ae. albopictus*, exhibited BFV-positive saliva across all four temperature profiles (18°C, 21°C, 24°C or 27°C) examined. These two invasive mosquito species must therefore be classified as potential vectors for BFV, indicating the potential risk of BFV transmission outside of Australia.

## Introduction

Human infections with arboviruses (arthropod-borne viruses) like dengue and chikungunya virus (CHIKV) have risen globally in recent decades, driven by climate change, land-use shifts, and globalization, which support both the spread of invasive mosquitoes/virus introduction into new regions and the subsequent emergence (1). One important family of the arboviruses is the family of *Togaviridae*, and more specific the genus of *Alphaviruses*, where also CHIKV belongs to. However, another member of this genus, is Barmah Forest virus (BFV), first isolated in 1974 from *Culex annulirostris* mosquitoes in Australia collected in the Barmah State Forest in Northern Victoria (2) and in the same year from *Cx. annulirostris* and *Aedes normanensis* in Queensland ((3), identified as BFV in (4)). It has been associated with human disease since 1988 (5). Its clinical presentation is similar to other alphavirus infections such as Ross River virus (RRV) (6), CHIKV, O’nyong-nyong virus, and Mayaro virus (7). Typical symptoms include fever, arthritis, headache, rash, joint swelling, and pain (6,8). After RRV, BFV is the second most common arbovirus infection in Australia, causing several 100 cases annually and over 16,000 confirmed cases since 2000 (9). However, it can be assumed that the actual number of BFV infections is much higher, as misdiagnoses as CHIKV or RRV are considered common (10). Large outbreaks of BFV occurred in the Northern Territory in 1992 (11), in Western Australia in 1993–1994 (12), in New South Wales in 1994–1995 (13), in Victoria in 2002 (14), and in Southeastern Queensland in 2002–2003 (15). Initially, BFV was considered to be limited to Australia, but in 2014, a case was reported in a child from Papua New Guinea with no travel history (16). Genetic analyses suggest an introduction in the early 1900s by humans, livestock or mosquitoes from Australia. In 2019, the first outbreak of BFV in Tasmania was reported (17). Since then, it has been observed in all states of mainland Australia, in Papua New Guinea and in Tasmania (10, 17).

Animals susceptible to BFV are not well defined, but it is assumed that BFV can infect a broader range of vertebrate hosts compared to RRV (18). Seroprevalence studies revealed BFV antibodies in different marsupials and other nonhuman vertebrates, like horses, brushtail possums, cats and dogs (19-21). The detection of antibodies also suggests that migrating hosts like birds might be part of the transmission cycle (22) and humans might act as reservoir hosts (23). So far, no symptomatic BFV infections in non-human hosts are known (21).

Based on isolations of BFV from mosquitoes collected in the field, BFV might has a broad spectrum of mosquito vectors. BFV has been isolated from different field collected mosquito species: *Ae. camptorhynchus* (12,24), *Ae. normanensis* (25), *Ae. vigilax* (13,25,26), *Cx. annulirostris* (25,27), *Cx. quinquefasciatus* (27), *Coquillettidia xanthogaster* (27), *Cq. linealis* (26), *Anopheles annulipes* s.l. (28). In addition, BFV was detected in the biting midge species *Culicoides marksi* (29). However, it should be noted that the detection of BFV in vectors collected in the field does not mean that these species are able to transmit the virus, since even a single blood meal from an infected host can lead to a positive result. Experimental studies are needed to confirm the ability of the species to transmit BFV.

Under laboratory conditions, vector competence has been demonstrated for three *Aedes* species: *Ae. notoscriptus* (30,31), *Ae. vigilax* (30,32,33), *and Ae. procax* (33), but not for *Ae. aegypti* (34). In addition, vector competence was shown for *Verrallina funerala* (35) and *Coquillettidia linealis* (26). *Culex annulirostris* is able to transmit BFV on a low level (transmission rates > 10%), whereas *Cx. quinquefasciatus and Cx. sitiens* should not be considered as vectors (34). Transmission of BFV by *Ae. vigilax* has been shown as early as 3 days post infection and by *Cq. linealis* as early as 6 days post infection (26,32). It is assumed, that *Ae. vigilax* (13,15,25,32, 33,36) and *Ae. camptorhynchus* (24), although there are no vector competence studies for *Ae. camptorhynchus*, are key vectors of BFV, which are both associated with coastal wetlands. These species have in common, that they are all restricted to the Australasian region.

Understanding which mosquitoes can transmit BFV is crucial. Since travel and trade increase, the risk of spreading BFV to other continents is potentially high. Climate change may also influence mosquito distribution and virus transmission (23,36). The related alphavirus CHIKV, serves as a cautionary example: it has spread across all continents in the last decades, causing millions of infections annually (37). This study aims to analyze a wider range of mosquito species for their vector competence for BFV transmission beyond Australasian species, to better assess the potential risk of introduction and subsequent spread at a global scale. Therefore, we included *Culex* and *Aedes* species, as Australian mosquito species of both genera show transmission potential for BFV (32,34). We included *Cx. quinquefasciatus*, a species widespread in tropical regions, and *Cx. torrentium*, which is native to Europe and demonstrated to have a high vector competence for a range of arboviruses including Sindbis virus as an Alphavirus (38-41). We also examined the invasive species *Ae. aegypti* and *Ae. albopictus*, two globally distributed species that play a central role in chikungunya virus transmission (42) As there have been no laboratory studies on the vector competence of *Anopheles* mosquitoes for BFV, we included *An. stephensi* in our investigations, an invasive mosquito species spreading within Asia and Africa (43).

## Materials and Methods

### Collecting and rearing of mosquitoes

During summer 2024, egg rafts of *Cx. torrentium* were collected in the field in northern Germany (Lon: 53.467821/Lat: 9.831346). Eggs and larvae were kept at room temperature under a 12:12 light:dark photoperiod. Species identification was conducted by isolating DNA from a pool of 5 L1/L2 larvae per egg raft (DNeasy blood & tissue kit, Qiagen, Hilden, Germany) followed by a multiplex quantitative real-time PCR (qPCR) (44). Pupae and adults were housed in an insectary with a relative humidity of 70%, 26°C, and a 12:12 light:dark photoperiod, incorporating a 30 min twilight. To eliminate the possibility of natural arbovirus infections, 10 adult mosquitoes were randomly chosen and tested by pan-Orthobunya-, pan-Flavivirus-, and pan-Alphavirus-PCR (45-47). All screened specimens tested negative. Laboratory colonies of *Cx. quinquefasciatus, Ae. aegypti* (both long-established colonies received by Bayer, Leverkusen, Germany), and *Ae. albopictus* (established from eggs collected in the field (Heidelberg, Germany) in 2016/2017) were reared under the same insectary conditions. *Anopheles stephensi* (strain SxK Nijmegen) were reared in an insectary with a relative humidity of 70%, 28°C, and a 12:12 light:dark photoperiod, incorporating a 30 min twilight.

### Infection of mosquitoes

Female mosquitoes in an age between 4 to 14 days were deprived of food prior the infection experiments. While most species were starved for 24 hours, *Cx. torrentium* underwent an extended starvation period of 48 hours. An artificial blood meal was offered for two hours at 24°C, containing 50% human blood (type 0, expired transfusion reserves), 30% of an 8% fructose solution, 10% filtrated bovine serum (FBS), and 10% virus stock, final virus concentration was 6.88 × 10^7^ FFU/mL. The BFV stock (strain BH2193 / U73745) was propagated in Vero cells (*Chlorocebus sabaeus*, CVCL_0059, obtained from ATCC, cat # CCL-81). An artificial blood meal was offered either via cotton stick (*Culex*) or via two 50µL drops on the bottom of the vial (*Aedes, Anopheles*). Feeding rates (FR, number of engorged females per number of fed females) are shown in table 1. Only fully engorged specimens were incubated for 7 days at 70% humidity and temperature profiles of a daily average of 18°C, 21°C, 24°C or 27°C with a fluctuation of ±5°C within 24 h. To simulate natural day-night temperature cycles, peak temperatures were programmed to occur midway through the light phase, while minimum temperatures coincided with the midpoint of the dark phase. Mosquitoes had continuous access to fructose solution provided on a cotton pad, which was renewed every 2-3 days. The survival rate (SR, number of alive mosquitoes 7 days post infection per number of engorged females) of all mosquito species are shown in table 1. All experiments were performed in 2 replicates.

**Table 1.**
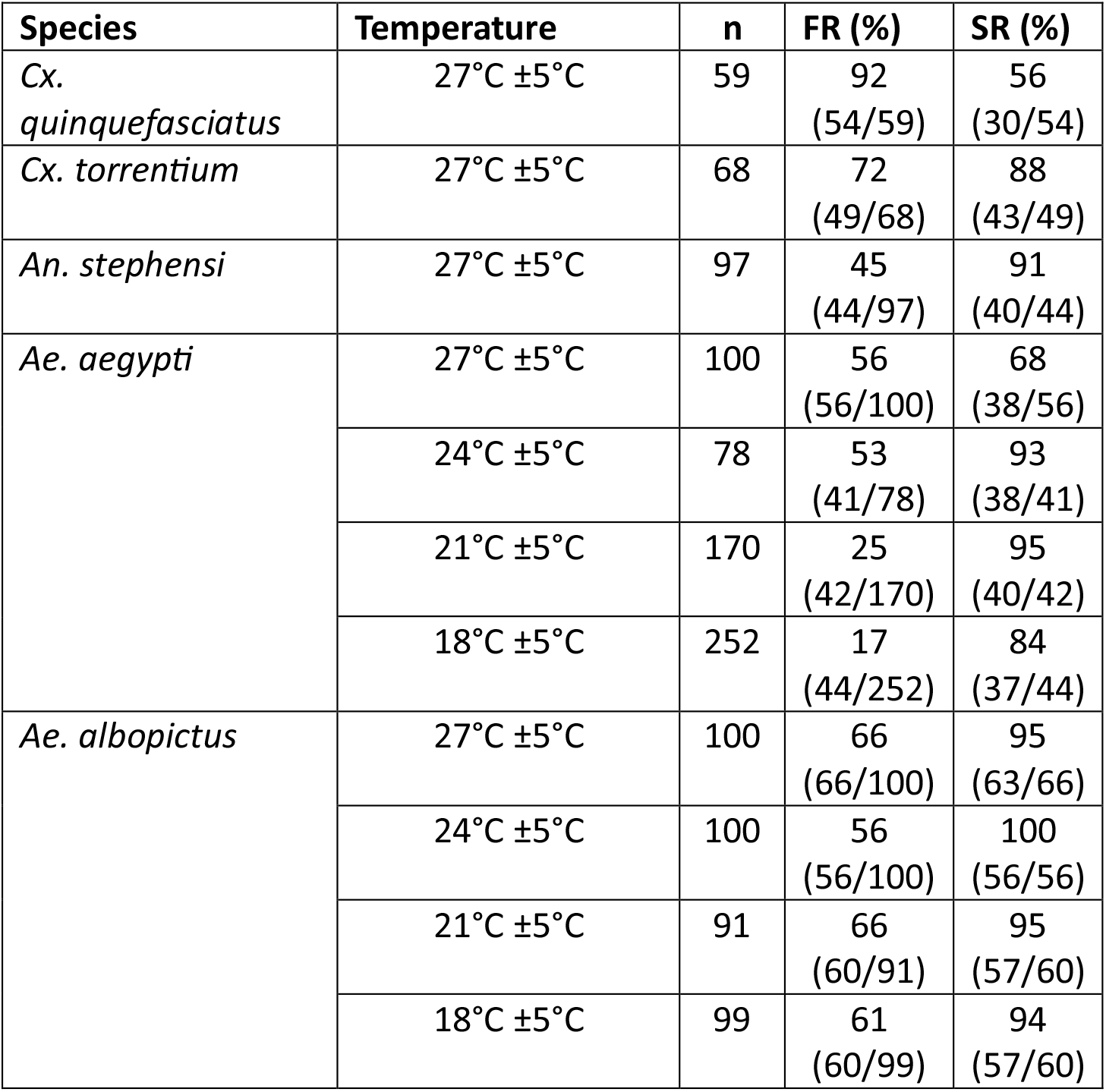
Feeding- and survival rates of all investigated species. n = Total input of mosquitoes per experiment condition, FR = feeding rate, number of engorged females per total input (n), SR = survival rate, number of alive mosquitoes seven days post infection per number of engorged females

| Species | Temperature | n | FR (%) | SR (%) |
| --- | --- | --- | --- | --- |
| *Cx. quinquefasciatus* | 27°C ±5°C | 59 | 92<br>(54/59) | 56<br>(30/54) |
| *Cx. torrentium* | 27°C ±5°C | 68 | 72<br>(49/68) | 88<br>(43/49) |
| *An. stephensi* | 27°C ±5°C | 97 | 45<br>(44/97) | 91<br>(40/44) |
| *Ae. aegypti* | 27°C ±5°C | 100 | 56<br>(56/100) | 68<br>(38/56) |
|  | 24°C ±5°C | 78 | 53<br>(41/78) | 93<br>(38/41) |
|  | 21°C ±5°C | 170 | 25<br>(42/170) | 95<br>(40/42) |
|  | 18°C ±5°C | 252 | 17<br>(44/252) | 84<br>(37/44) |
| *Ae. albopictus* | 27°C ±5°C | 100 | 66<br>(66/100) | 95<br>(63/66) |
|  | 24°C ±5°C | 100 | 56<br>(56/100) | 100<br>(56/56) |
|  | 21°C ±5°C | 91 | 66<br>(60/91) | 95<br>(57/60) |
|  | 18°C ±5°C | 99 | 61<br>(60/99) | 94<br>(57/60) |

### Analysis

To evaluate the transmission rate (TR, number of BFV-positive saliva per number of positive bodies) and the transmission efficiency (TE, number of BFV-positive saliva per number of fed females), a salivation assay was conducted as previously described (48). In short, saliva from each specimen was collected for 30 minutes in a tip containing 10 µL phosphate-bufferd saline (PBS). The saliva solution was applied on Vero cells and incubated at 37°C and 5%-CO_2_ for five days. If cells showed a cytopathic effect, the supernatant was harvested, RNA was extracted and a qPCR was performed to confirm BFV infection. Reverse transcription (RT) qPCR was performed after the protocol of Inglis et al. (27) using the QuantiTect Reverse Transcription Kit (Qiagen, Hilden, Germany), with an addition of the VetMAX^™^ Xeno^™^ Internal Positive Control (Applied Biosystems, Thermo Fisher Scientific Corporation, Waltham, MA, USA). For quantification, a standard was designed (ATGCCTCAGGGCCACTCACCATACTTGTGGTAGCTATTATAGTCGTTGTTGTAGTATCCATTGTAGTATGTGCAAGACACTAGCAGAACTA).

The RT-qPCR was validated following the guidelines outlined by Bustin *et al*. in their publication “Minimum Information for Publication of Quantitative Real-Time PCR Experiments” (49). Ten-fold serial dilutions of the BFV standard, ranging from 1.09 to 1.09 × 10^11^ copies/µL of the above-mentioned standard were analyzed in five replicates using the previously described RT-qPCR protocol. The limit of detection was identified as 1.09 × 10^2^ copies/reaction, or 1.23 × 10^4^ copies/mosquito body, with a standard deviation of 0.527 C_q_s at this concentration. The linear dynamic range extended from the detection limit to ≥1.23 × 10^13^ copies/mosquito body, proving a proportional relationship between concentration and PCR signal within this range, ensuring reliable results. The calibration curve resulted in a coefficient of determination of 0.993, a slope of -3.772, a y-intercept of 45.112, and a PCR efficiency of 0.841, i.e. 84.1% of the target molecules are successfully amplified in each cycle.

Additionally, RNA of mosquito bodies (including the head) was extracted and qPCR was performed, to calculate the infection rate (IR, number of BFV-positive bodies per fed females) and the mean body virus titer.

## Results

At 7 days post infection no BFV positive saliva was detected for *Cx. quinquefasciatus* and *Cx. torrentium* (table 2). The infection rate (IR) in these species was 12.9% and 23.3%, respectively, with a mean body titer of 4.6 and 4.8 log10 BFV RNA copies per specimen (table 2). Similarly, *An. stephensi* showed an IR of 16.7% with a mean body titer of 5.0 log10 BFV RNA copies per specimen and no transmission (table 2).

**Table 2.** Infection rate, transmission rate and transmission efficiency of different mosquito species for BFV. dpi = days post infection, n = Number of fed and investigated specimens, IR = infection rate, number of BFV-positive specimens per engorged specimens, TR = transmission rate, number of BFV-positive saliva per BFV-positive bodies, TE = transmission efficiency, number of BFV-positive saliva per number of engorged females

| Species | dpi | Temperature | n | IR (%) | TR (%) | Mean body titer<br>log<sub>10</sub> RNA<br>copies/mosquito<br>specimen (95%<br>confidence<br>interval) | TE (%) |
| --- | --- | --- | --- | --- | --- | --- | --- |
| *Cx. quinquefasciatus* | 7 | 27°C ±5°C | 30 | 23.3<br>(7/30) | 0.0<br>(0/7) | 4.6 (4.2 – 5.0) | 0.0<br>(0/30) |
| *Cx. torrentium* | 7 | 27°C ±5°C | 31 | 12.9<br>(4/31) | 0.0<br>(0/4) | 4.8 (3.8 – 5.9) | 0.0<br>(0/31) |
| *An. stephensi* | 7 | 27°C ±5°C | 30 | 16.7<br>(5/30) | 0.0<br>(0/5) | 5.0<br>(2.3 – 7.8) | 0.0<br>(0/30) |
| *Ae. aegypti* | 7 | 27°C ±5°C | 30 | 70.0<br>(21/30) | 28.6<br>(6/21) | 6.3<br>(5.3 – 7.2) | 20.0<br>(6/30) |
|  | 7 | 24°C ±5°C | 30 | 60.0<br>(18/30) | 16.7<br>(3/18) | 6.5<br>(5.3 – 7.6) | 10.0<br>(3/30) |
|  | 7 | 21°C ±5°C | 30 | 80.0<br>(24/30) | 25.0<br>(6/24) | 6.1<br>(5.1 – 7.0) | 20.0<br>(6/30) |
|  | 7 | 18°C ±5°C | 33 | 63.6<br>(21/33) | 28.6<br>(6/21) | 5.2<br>(4.4 – 6.0) | 18.2<br>(6/33) |
| *Ae. albopictus* | 7 | 27°C ±5°C | 32 | 65.6<br>(21/32) | 47.6<br>(10/21) | 5.8<br>(4.7 – 6.8) | 31.3<br>(10/32) |
|  | 7 | 24°C ±5°C | 30 | 66.7<br>(20/30) | 25.0<br>(5/20) | 5.3<br>(4.6 – 6.0) | 16.7<br>(5/30) |
|  | 7 | 21°C ±5°C | 29 | 72.4<br>(21/29) | 4.8<br>(1/21) | 5.3<br>(4.6 – 5.9) | 3.5<br>(1/29) |
|  | 7 | 18°C ±5°C | 29 | 75.9<br>(22/29) | 4.5<br>(1/22) | 5.5<br>(4.8 – 6.3) | 3.4<br>(1/29) |

In contrast, both *Ae. aegypti* and *Ae. albopictus* showed positive saliva at all four investigated temperature profiles (table2). For *Ae. aegypti*, we detected IRs from 60% to 80% with a mean RNA body titer between 5.2 and 6.5 log10 BFV copies per specimen (table2). TRs for *Ae. aegypti* ranged from 16.7% to 28.6% with TEs between 10% and 20% (table2), without major differences between the analyzed temperature profiles. *Aedes albopictus* showed comparable IRs, ranging from 65.6% to 75.9%, with slightly lower mean viral loads, ranging from 5.3 to 5.8 log10 BFV copies per specimen (table2). Transmission values in *Ae. albopictus* were notably higher than in *Ae. aegypti* at 27°C, showing a TR of 47.6% and a TE of 31.3% (table2). At 24°C, with a TR of 25% and a TE of 16.7%, *Ae. albopictus* showed values in the same range as *Ae. aegypti*. In contrast to *Ae. aegypti*, a clear trend of lower transmission at lower temperatures of 21°C and 18°C were observed for *Ae. albopictus*, showing TRs of 4.8% and 4.5% and TEs of 3.5% and 3.4%.

## Discussion

Our results demonstrate that *Culex, Anopheles*, and *Aedes* species are susceptible to BFV infection under laboratory conditions. However, only the two tested *Aedes* species, *Ae. aegypti* and *Ae. albopictus*, were capable of transmitting the virus, as indicated by the presence of infectious virus particles in their saliva. To our knowledge, this is the first time transmission of BFV by these species has been shown. *Aedes albopictus* was not tested before, but the observations for *Ae. aegypti* are in contrast to studies by Boyd et al., who did not observe transmission of BFV by this species, even though infection was detected (34). This discrepancy can likely be explained by the lower input titer used in their study. To the best of our knowledge, there is no information available on the viraemia in hosts or on the minimum infectious dose required for the transmission of BFV to mosquitoes, further research is needed to fill this knowledge gap.

So far, no studies on the vector competence of *An. stephensi* and *Cx. torrentium* for BFV have been published. *Culex quinquefasciatus* has been investigated previously, and in line with our results, no transmission of BFV was observed (34). In fact, the only investigated *Culex* species that showed low transmission potential is *Cx. annulirostris*. However, there are also studies that did not detect any transmission of BFV for this species (33,34). This suggests that vector competence for BFV may be largely restricted to *Aedes* species, despite a broader infection susceptibility across other mosquito genera.

These findings are particularly relevant for risk assessment of BFV transmission in Australia, where *Ae. aegypti* is present and plays role as vector of dengue virus in some regions (50) but has not been considered a potential vector of BFV so far (34). BFV transmission has mainly been linked to coastal Australian *Aedes* species such as *Ae. camptorhynchus* and *Ae. vigilax* (24,36). As *Ae. aegypti* is distributed across all continents except Antarctica (51), BFV transmission could potentially occur outside Australia. In addition, *Ae. albopictus* is not yet established on the Australian mainland, but repeated introductions have been reported in the Torres Strait, and BFV transmission has already occurred in neighbouring Papua New Guinea (52). Given it’s demonstrated vector competence for BFV, the establishment of *Ae. albopictus* in Australia would increase the risk of transmission. Due to the global distribution of the species, our findings highlight the potential risk for BFV emergence in other parts of the world. In Europe, *Ae. albopictus* has been linked to all known autochthonous outbreaks of CHIKV, as well as dengue and Zika virus infections (53). This highlights the particular relevance of *Ae. albopictus* as a competent vector. Interestingly, both CHIKV and BFV can be transmitted at relatively low temperatures (15°C ±5°C and 18°C ±5°C, respectively) (54), which are typical for temperate regions such as Central Europe.

Another interesting aspect is, that BFV most likely has a broader spectrum of potential hosts compared to other alphaviruses (18,34), although recent studies suggest a broader host range for CHIKV and RRV as well (55,56). While RRV and CHIKV transmission is largely associated with specific marsupial or primate hosts, respectively (57), BFV has been detected in multiple vertebrate species, including birds and various mammals (19-22). This broader host range may increase the likelihood of BFV establishing sustained enzootic transmission cycles in new environments and thereby facilitate geographic expansion, but a full risk assessment would require a better understanding of the transmission cycle of BFV and especially the clear identification of amplification and reservoir hosts.

## Conclusion

This study provides novel insights into the mosquito vector of BFV, a topic that remains to a large extent underexplored, especially for mosquito species not native to Australia. We could show, that the two invasive mosquito species *Ae. albopictus* and *Ae. aegypti* are potent vectors for BFV. Given the potential for arbovirus emergence in new regions due to globalization and climate-driven shifts in vector distributions, our results indicating the potential risk of BFV transmission outside of Australia.

### Statement

Large language models (LLMs) were used during the preparation of this manuscript to support language editing and improve clarity and readability. The authors reviewed, verified, and take full responsibility for the final content of the manuscript. The use of LLMs did not replace the authors’ scientific judgment, interpretation of results, or responsibility for the work.

## Conceptualization

A.H., S.J.; Investigation: A.H., P.H., U.L., A.M., S.J.; Methology: A.H., P.H., S.J.; Formal analysis: A.H.; Original draft preparation: A.H., S.J.; Review and editing: all authors; Funding acquisition: J.S.C., A.H., R.L.; Resources: R.L., N.B.; Supervision: A.H., S.J.; All authors have read and agreed to the published version of the manuscript.

